# The immediate early gene *Arc* is not required for hippocampal long-term potentiation

**DOI:** 10.1101/2020.01.03.893305

**Authors:** M. Kyrke-Smith, L.J. Volk, S.F. Cooke, M.F. Bear, R.L. Huganir, J.D Shepherd

**Affiliations:** Department of Neurobiology and Anatomy, University of Utah; Department of Neuroscience, The Johns Hopkins School of Medicine; Department of Neuroscience, University of Texas Southwestern; The Picower Institute of Learning and Memory, MIT; Department of Basic and Clinical Neurosciences, Kings College London (KCL); MRC Centre for Neurodevelopmental Disorders, KCL

## Abstract

The immediate early gene *Arc* is critical for maintenance of long-term memory. How Arc mediates this process remains unclear, but it has been proposed to sustain Hebbian synaptic potentiation, which is a key component of memory encoding. This form of plasticity is modelled experimentally by induction of long-term potentiation (LTP), which increases Arc mRNA and protein expression. However, mechanistic data implicates Arc in the endocytosis of AMPA-type glutamate receptors and the weakening of synapses. Here, we took a comprehensive approach to determine if Arc is necessary for hippocampal LTP. We find that Arc is not required for LTP maintenance and must regulate memory storage through alternative mechanisms.

## INTRODUCTION

The maintenance of memory and Hebbian forms of synaptic plasticity, such as long-term potentiation (LTP) and depression (LTD), requires activity-dependent gene expression and de novo protein synthesis (Goelet et al., 1986; Bliss and Collingridge, 1993; Abraham and Williams, 2003; Takeuchi et al., 2013; Poo et al., 2016; Choi et al., 2018; Kyrke-Smith and Williams, 2018). The molecular underpinnings of LTP, in particular, are of immense interest due to the considerable overlap in activity-induced genes critically involved in both LTP and memory (Alberini, 1999, 2009; Kandel et al., 2014). Thus, identifying genes that are necessary for the maintenance of LTP may be key to identifying ‘memory molecules’. The immediate early gene *Arc* is induced by neuronal activity, is necessary for memory consolidation (Guzowski et al., 2000; Plath et al., 2006; Ploski et al., 2008), and implicated in various forms of synaptic plasticity (Shepherd and Bear, 2011). *Arc* mRNA expression is robustly induced in the dentate gyrus of the hippocampus after seizure or LTP (Link et al., 1995; Lyford et al., 1995). The newly expressed *Arc* mRNA moves out of the cell body and into dendrites where it specifically localises near potentiated synapses (Steward et al., 1998; Yin et al., 2002; Moga et al., 2004; Rodriguez et al., 2005). LTP maintenance has been shown to be attenuated by application of antisense RNA against Arc (Guzowski et al., 2000; Messaoudi et al., 2007) and in Arc KO animals under urethane anaesthesia (Plath et al., 2006). However, Arc is also critical for the maintenance of hippocampal LTD (Plath et al., 2006; Waung et al., 2008) and for homeostatic scaling of AMPA-type glutamate receptors (AMPARs) (Shepherd et al., 2006; Korb et al., 2013). How one protein can be critical for both increases and decreases in synaptic strength has remained an open question.

Arc protein interacts with the endocytic machinery at synapses, such as clathrin-adaptor protein 2 (AP-2), (DaSilva et al., 2016) dynamin and endophilin (Chowdhury et al., 2006a). Acute over expression of Arc leads to the endocytosis of AMPARs, thereby decreasing synaptic strength (Rial Verde et al., 2006; Shepherd et al., 2006). Additionally, Arc interacts with the calcium/calmodulin-dependent protein kinase II– beta (CaMKIIβ) isoform to decrease synaptic strength at inactive synapses (Okuno et al., 2012). However, Arc protein can also interact with drebrin A, which may lead to F-Actin stabilization and the enlargement of dendritic spines, indicative of increased synaptic strength (Messaoudi et al., 2007; Bramham et al., 2010; Nair et al., 2017). The multitude of Arc protein-protein interactions has made it difficult to determine how Arc mediates long-term memory, and raises the possibility that Arc may facilitate memory through mechanisms other than direct regulation of synaptic strength. For example, Arc protein may be found in the nucleus where it interacts with the nuclear spectrin isoform βSpIVΣ5 (Bloomer et al., 2007) and the histone acetyl-transferase Tip60 (Wee et al., 2014). Additionally, we recently found that Arc can form virus-like capsids that can encapsidate RNA and transfer Arc mRNA between cells (Pastuzyn et al., 2018).

Given the extensive literature implicating LTP as a cellular correlate of memory (refs), we aimed to comprehensively investigate whether Arc is directly required for the maintenance of hippocampal LTP. We used two different *Arc* knock-out (Arc KO) mouse lines and a conditional KO floxed line (Arc cKO), for both *in vitro* and *in vivo* LTP analysis. We find that Arc is not required for the maintenance of LTP, using multiple LTP induction protocols. These findings suggest that Arc’s role in long-term memory consolidation is most likely mediated through mechanisms that do not involve the maintenance of synaptic potentiation.

## MATERIALS AND METHODS

### Animals

Experiments were performed in three different Arc KO models. Two full Arc knockout animal lines were used: the Arc^tm1Stl^ (Wang et al., 2006) line, where GFP was knocked in to the endogenous Arc locus (Arc^GFP/GFP^) and the Kuhl Arc^−/−^ line (Plath et al., 2006). An Arc conditional KO line (Arc^Fl/Fl^) that was previously generated (provided, by Dr. Richard Palmiter, University of Washington) was also used. In this line, loxP sites were inserted to enable the removal of the entire Arc gene upon introduction of cre-recombinase (Chen et al., 2018). All lines are on the C57BL6 background and WT littermates were used as controls. All procedures were approved by the Institutional Animal Care and Use Committees of the University of Utah, The Johns Hopkins University, and the Massachusetts Institute of Technology, in conjunction with NIH guidelines.

### In vitro electrophysiology

Slices were prepared from WT and KO mice at age 6 – 8 weeks (Arc cKO and Arc^GFP/GFP^) or 11 – 14 weeks (Arc^−/−^). Mice were anesthetized with the inhalation anesthetic Isoflurane prior to decapitation. 400μm transverse slices were prepared using a Leica VT1200S vibratome (Arc^−/−^) or a Leica VT1000S vibratome (Arc^GFP/GFP^) in ice cold oxygenated (95% O_2_/5% CO_2_) dissection buffer containing the following (in mM): 2.6 KCl, 1.25 NaH_2_PO_4_, 26 NaHCO_3_, 211 sucrose, 10 glucose, 0.75 CaCl_2_, 7MgCl_2_. Slices were recovered in a static submersion chamber for at least 2hr in 30°C aCSF containing the following (in mM): 125 NaCl, 3.25 KCl (5 KCl_Arc^−/−^), 25 NaHCO_3_, 1.25 NaH2PO_4_·H2O, 11 glucose, 2CaCl_2_, 1MgCl_2_. Field excitatory postsynaptic potentials (fEPSPs) were evoked at 0.033Hz with a 125mm platinum/ iridium concentric bipolar electrode (FHC, Bowdoinham, ME; Arc^−/−^) or 75 mm Tungsten bipolar electrode (World Precision Instruments, Sarasota, FL; Arc^GFP/GFP^) placed in the middle of stratum radiatum of CA1. A 1-4 MΩ glass recording electrode filled with aCSF was positioned 200 - 400 μm away (orthodromic) from the stimulating electrode. Input-output (I/O) curves were obtained for each slice and stimulus intensity for all subsequent recordings was set to elicit a fEPSP slope that was 30 - 40% of the maximum response. Recording aCSF and temperature were identical to recovery conditions, with a flow rate of ~3mL/min. Stimulation protocols were as follows: <u>E-LTP:</u> 20 pulses at 100 Hz. <u>Moderate Frequency stimulation:</u> 900 pulses at 10 Hz. <u>High frequency stimulation LTP:</u> 2 trains of 100 pulses at 100 Hz. <u>Theta burst L-LTP:</u> 4 trains at 10 second inter-burst interval. Each train consisted of 10 bursts at 5Hz, with each burst containing 4 stimuli at 100Hz.

### In vivo electrophysiology

Mice were anesthetized under isoflurane (3% for induction and 1.5% for maintenance) and head-fixed in a stereotaxic frame. Rectal temperature was maintained at 37°C using a heat blanket. A dental drill was used to make craniotomies in the skull and electrodes were placed according to established stereotaxic coordinates on the left-hand side of the brain. For LTP experiments in the dentate gyrus, a concentric bipolar stimulating electrode (Rhodes Medical Instruments, California) was positioned in the medial perforant path (MPP), 3 mm left of lambda and at a depth of ~1.5 mm from brain surface. A glass micropipette was lowered into the hilus of the left dentate gyrus, 2 mm posterior to bregma, 1.6 mm left of the mid-line and at a depth of ~1.5 mm, to record positive-going evoked field responses next to the granule cell bodies. Correct placement of the stimulating electrode in the MPP was confirmed by the characteristic short latency onset of the evoked field response (2–2.5 msec) and the early onset of the population spike (4 msec approx.). For all experiments, input–output relationships were assessed using a range of stimulus intensities from 0-220 μA, which produced a close to maximal response. Five responses were collected at each intensity and averaged. Test responses for LTP experiments were evoked by monophasic stimuli set at an intensity to evoke a response ~50% of maximum fEPSP slope and a population spike of approximately 1 mV (100–220 μA, 60 μsec). A stable 30-minute baseline was acquired before delivering a theta burst tetanus to induce LTP. The tetanus comprised six series of six trains of six stimuli at 400 Hz with 200 msec between trains and 20 sec between series. Pulse width was doubled during the tetanus to 120 μsec. Only the very early component of the EPSP slope was measured for analysis in order to ensure that there was no contamination by the population spike. The slope of the fEPSP was expressed as a percentage change relative to the averaged baseline response. At the end of each experiment the anesthetized mouse was killed by cervical dislocation.

### Viral injection

Mice were anesthetized with isoflurane (3% for induction, 1.5-2% for surgery) and placed in a stereotaxic frame following the suppression of reflexes. The antibiotic Baytril (8 mg/kg, VetOne, Boise, ID), the analgesic Carprofen (5 mg/kg, Zoetis, Parsippany-Troy Hills, NJ) and the anti-inflammatory dexamethasone (13 mg/kg, VetOne, Boise, ID) were administered prior to placing the mouse in the stereotax to control pain/swelling. Lidocaine (100 mg/kg, VetOne, Boise, ID) was then injected subcutaneously beneath the scalp prior to incision. The skin was incised to expose the skull and a small “burr hole” was made with a pneumatic dental drill above the CA1 region of the hippocampus (350 μm lateral and 50 μm anterior to LAMBDA). A pulled glass pipette, backfilled with mineral oil and filled with virus, was lowered (250 μm below surface) and allowed to rest for 5 minutes. Bilateral injections of either AAV:CaMKII-GFP (3.18 × 10^8^ particles; Ed Boyden, University of North Carolina Vector Core) or AAV:CaMKII-GFP-cre (2.1 × 10^8^ particles; University of North Carolina Vector Core) were delivered using a Nanoject II Auto-Nanoliter Injector (Drummond Scientific, Broomall, PA). Following a further 5 min resting post-injection, the pipette was slowly removed. The scalp was then sutured closed and the animal was placed in a warm cage for recovery. Animals were maintained at ~37 °C throughout the procedure and recovery, and general condition and reflex signs were monitored closely. Mice were monitored postoperatively for signs of infection or discomfort. Animals recovered for 2 weeks to allow for viral expression before slice preparation. To quantify the effectiveness of cre to knockout Arc expression in these animals, slices used for electrophysiology were subsequently prepared for western blotting to assess Arc protein levels. Slices were only used for experiments if GFP could be detected along the entire CA1 region of each slice prior to electrophysiology. GFP was visualized in slices using an LED light source at 470 nm (pE-100, CoolLED, United Kingdom) on a SliceScope Pro mounted FireWire (B FWCAM X M) camera linked to the SCIlight 2.2 software (Scientifica, United Kingdom).

### Immunohistochemistry

Representative image of GFP injection was made from animal injected with AAV:CaMKII-GFP, as described above. After 2 weeks recovery, animal was perfused with 4% paraformaldehyde, followed by 24 h fixation in further 4% paraformaldehyde. The brain was cryoprotected in 30% sucrose before being sectioned (30 μm) on a cryostat. Sections were stained for DAPI (Thermo Fisher R37606, 5 min at room temperature) before mounting coverslips on slides in Fluoromount (Sigma F4680) and dried. Images were captured on the confocal microscope (Ti2, Nikon, Japan).

### Western blotting

After the CA1 region of the hippocampus was dissected from slices post-electrophysiology, tissue was mixed in 4 × Laemlli Buffer (40% glycerol, 250 mM Tris, 4% SDS, 50 mM STT, pH 6.8) and boiled at 95 °C for 10 min. SDS-PAGE gel electrophoresis was used to separate protein samples. Separated samples were transferred to a nitro-cellulose membrane (GE Healthcare). Following transfer, membranes were briefly stained with 0.1% Ponceau stain, destained with 1% acetic acid to remove background and then imaged to detect total protein. Membranes were blocked in 5% milk + 1 × tris-buffered saline (TBS; 10X: 152.3 mM Tris-HCL, 46.2 mM Tris base, 1.5 M NaCl, pH 7.6) for 30 min at room temperature. Membranes were then incubated with primary antibody against Arc (1:1000, custom rabbit polyclonal, Protein Tech) in 1 × TBS, overnight at 4 °C. Membranes were washed in 1 × TBS (3 × 10 min) before being incubated in a secondary antibody (1:10,000 HRP-conjugated goat anti-rabbit, Jackson ImmunoResearch), in blocking solution described above, for 1 h at room temperature. Membranes were then washed again in 1 × TBS (3 × 10 min). A chemiluminescent kit was used to detect protein bands (Bio-Rad, Hercules, CA). Membranes were imaged on an Azure c300 gel dock (Azure Biosystems, Dublin, CA). Blots were analysed and quantified using the Gel Analysis plugin in ImageJ.

### Statistics

For all LTP experiments, the magnitude of LTP over the first 5 min post-stimulation and the last 5 min of recording was averaged for each slice. Unpaired, two-tailed *t*-tests were performed on these averages from the Arc KO and WT littermates for each experiment. For I/O curves, linear regression was fitted for each curve (± 95% confidence interval) and an extra-sum-of-squares F test was used to compare slope of the best fit line between KO and WT groups. For paired pulse, a 2way ANOVA was performed to determine whether there was a genotype by ISI effect, with Sidak’s *post hoc* multiple comparison analysis of the difference between genotype at each ISI. For western blot analysis, Arc signal was normalised to total protein and then the fold change (FC) for all samples were calculated from the average of the GFP only group (sample/average of control group). Unpaired, two-tailed t-tests were then performed between the two groups. GraphPad Prism (GraphPad Software, San Diego, CA) was used for all statistics.

## RESULTS

### Arc is not required for LTP induced by high frequency stimulation in CA1

In order to comprehensively test the role of Arc in hippocampal LTP, we first used multiple high frequency stimulation (HFS) paradigms to induce plasticity at CA3 to CA1 synapses in hippocampal slices. Broadly, LTP can be divided into protein synthesis independent E-LTP or protein synthesis dependent L-LTP. As Arc expression increases within minutes of high frequency stimulation, we hypothesized that Arc would be required for the maintenance of LTP induced by HFS. To test this, we used a standard HFS protocol that typically induces LTP that persists for at least 1 hour. We used 6-8 week old Arc^GFP/GFP^ mice, which show no differences in basal transmission (WT slope = 2.237 ± 0.64, n = 20, Arc^GFP/GFP^ KO slope = 2.397 ± 0.80, n = 21; linear regression, extra-sum-of-squares F test between curves: F_(1,190)_ = 0.096, *p* = 0.756) (Figure 1A). There was no significant difference in the magnitude of LTP induced by a 2 × HFS (100 Hz) protocol between Arc^GFP/GFP^ KO and WT littermates (WT 178.84% ± 7.72, n = 5; KO 214.83% ± 24.44, n = 5; *t*-test*: p* = 0.20), nor was there a difference in the magnitude of LTP 1 hour post-stimulation (WT 146.48% ± 7.63, n = 5; Arc^GFP/GFP^ KO 141.29% ± 10.12, n = 5; *t*-test: *p* = 0.69) (Fig. 1B).

**Figure. 1.**
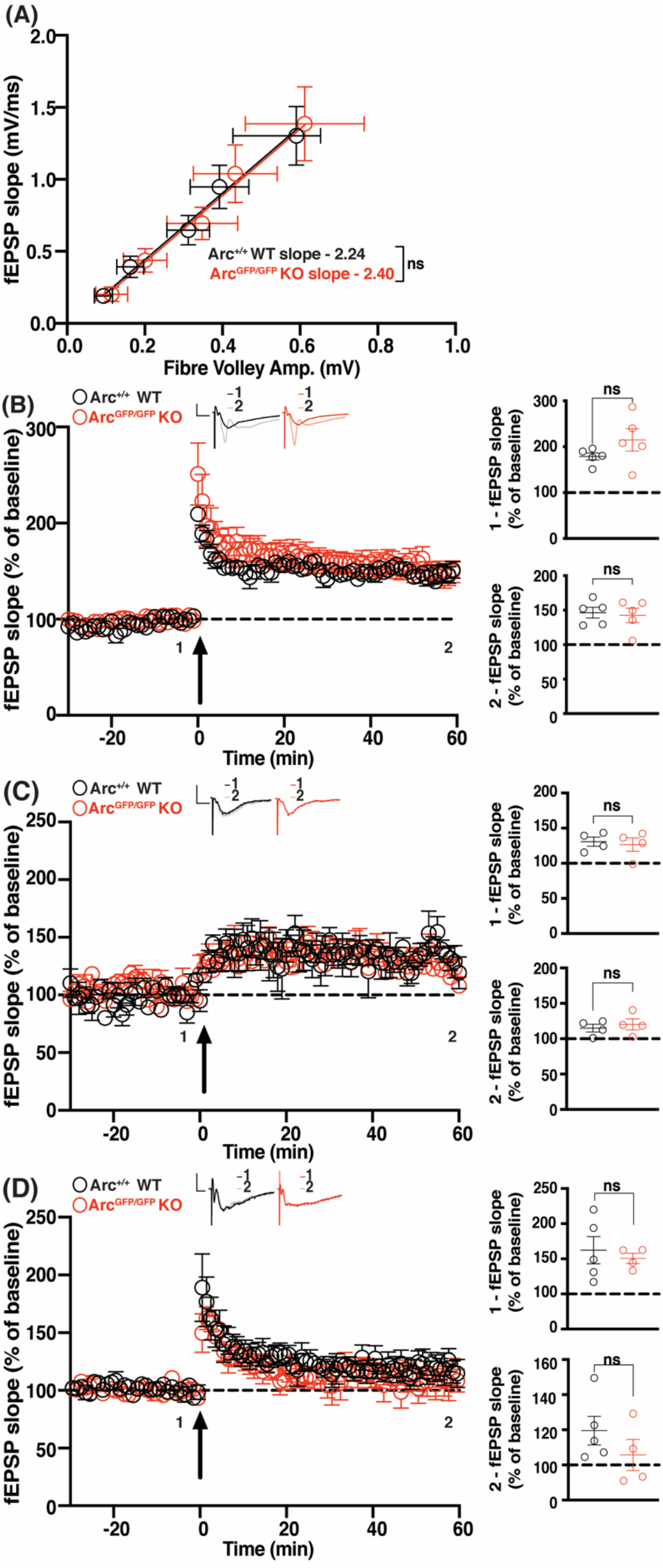
HFS CA1 LTP is normal in Arc^GFP/GFP^ KO animals. **(A)** There was no significant difference in basal synaptic transmission between Arc^GFP/GFP^ KO (red circles) and WT (black circles) littermates. **(B)** There was no significant difference in the average magnitude of LTP over the first 5 minutes post-stimulation (1), nor over the last 5 minutes of recording (2), between Arc^GFP/GFP^ KO and WT littermates when LTP was induced using a 2 × HFS (100 pulses at 100 Hz) protocol. **(C)** There was no significant difference in the magnitude of LTP over the first 5 min post-stimulation (1), nor over the last 5 minutes of recording (2), between the Arc^GFP/GFP^ KO and WT littermates when LTP was induced using a moderate frequency protocol (900 pulses at 10 Hz). **(D)** There was no significant difference in the magnitude of E-LTP over the first 5 min post-stimulation (1), nor over the last 5 minutes of recording (2), between the Arc^GFP/GFP^ KO and WT littermates when E-LTP was induced using weak, high frequency stimulation protocol (20 pulses at 100 Hz). Data are represented as mean ± SEM. Scale bars, 0.5 mV (horizontal) and 5 ms (vertical).

Surprised by this result, we hypothesised that the loss of Arc may lower the threshold for LTP induction (Shepherd and Bear, 2011). To determine this, we tested a moderate frequency stimulation protocol of 900 pulses at 10 Hz that typically induces little to no change in synaptic strength (Dudek and Bear, 1992). There was no significant difference in the magnitude of LTP induced between Arc^GFP/GFP^ KO and WT littermates (WT 124.7% ± 6.67, n = 4; Arc^GFP/GFP^ KO 125.3% ± 9.65, n = 4; *t*-test: *p* = 0.96), nor a difference in the magnitude of LTP 1 hour post-stimulation (WT 115% ± 5.35, n = 4; Arc^GFP/GFP^ KO 120.2% ± 7.74, n = 4; *p* = 0.60) (Fig. 1C). To further test the threshold hypothesis, we used a low repetition, high-frequency stimulation protocol that typically only induces E-LTP (20 pulses at 100 Hz) (Frey and Morris, 1997). There was no significant difference in the magnitude of LTP induced between Arc^GFP/GFP^ KO and WT littermates (WT 162.25% ± 19.4, n = 5, Arc^GFP/GFP^ KO 150.65% ± 7.21, n = 4; *t*-test: *p* = 0.63) nor was there a difference in the magnitude of LTP 1 hour post-stimulation (WT 119.54% ± 8.12, n = 5, Arc^GFP/GFP^ KO 105.6% ± 8.81, n = 4; *t*-test: *p* = 0.29) (Fig. 1D). These results show that Arc is not required for HFS induced LTP, nor does loss of Arc change the threshold of LTP induction in the CA1 region of the hippocampus.

### CA1 LTP induced by theta burst stimulation is increased in Arc KO mice

We tested another robust and commonly used L-LTP induction protocol, theta burst stimulation (TBS) (Larson et al., 1986; Thomas et al., 1998; Volk et al., 2013). We recorded for 2 hours post LTP induction to determine whether longer-term maintenance of L-LTP was affected. The initial magnitude of LTP induced was not significantly different between Arc^GFP/GFP^ KO and WT littermates (WT 222.0% ± 30.7, n = 6; KO 255.6% ± 34.3, n = 7; *t*-test: *p* = 0.48). However, 2 hours post-stimulation the magnitude of LTP was significantly greater in Arc^GFP/GFP^ KO mice compared with WT littermates (WT 132.9% ± 8.17, n = 6; KO 215.4% ± 14.46, n = 7; *t*-test: *p* = 0.0005. Fig. 2A). Since this Arc KO line expresses GFP from the Arc locus, we also investigated whether TBS L-LTP was altered in Arc^−/−^ mice (Plath et al., 2006). Interestingly, we find a small but significant difference in basal transmission in this line (WT slope = 4.97 ± 0.60, n = 16; Arc^−/−^ KO slope = 3.4 ± 0.36, n = 15; linear regression, extra-sum-of-squares F test between curves: F_(1, 175)_ = 21.37, *p* < 0.0001) (Fig. 2B). The initial magnitude of L-LTP induced was significantly greater in Arc^−/−^ KO mice compared to WT littermates (WT 230.46% ± 8.88, n = 10; KO 300.63% ± 14.45, n = 10; *t*-test: *p* = 0.0006). Similarly, the magnitude of LTP 2 hours post-induction was significantly greater in Arc^−/−^ KO mice than WT littermates (WT 155.11% ± 8.27, n = 10; KO 194.72% ± 8.55, n = 10; *t*-test: *p*=0.003) (Fig. 2C). These results show that Arc is not required for the maintenance of TBS L-LTP, and that loss of Arc may lead to an increase in TBS LTP magnitude.

**Figure. 2.**
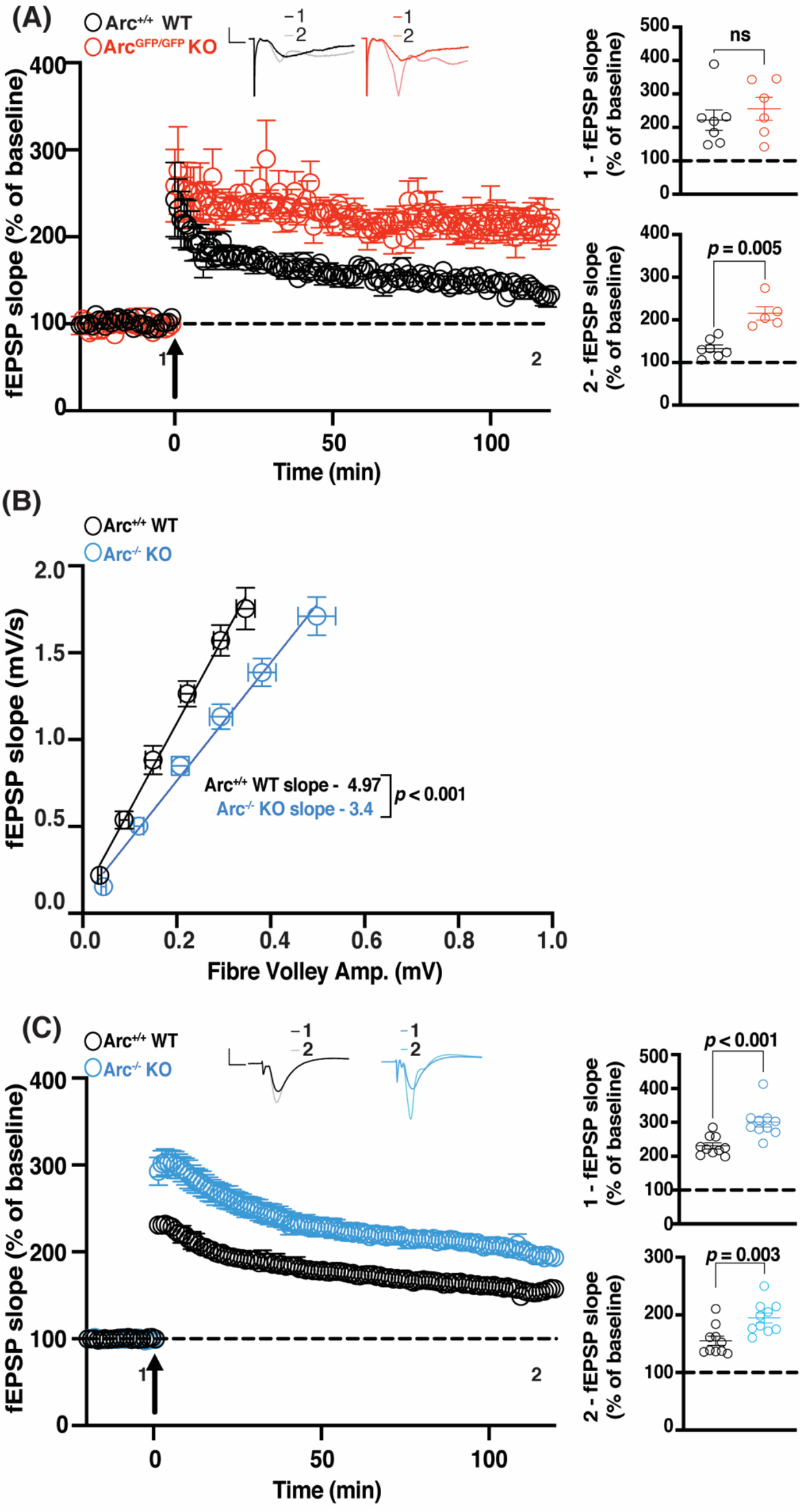
TBS CA1 LTP is enhanced in Arc KO animals. **(A)** There was no significant difference in the magnitude of LTP over the first 5 min post-stimulation (1) between the Arc^GFP/GFP^ KO (red circles) and WT (black circles) littermates, but the magnitude of LTP was significantly greater in Arc^GFP/GFP^ KO mice compared to WT littermates over the last 5 min of recording (2), when LTP was induced using a theta-burst stimulation protocol (4 trains at 10 second inter-burst interval. Each train consisted of 10 bursts at 5Hz, with each burst containing 4 stimuli at 100Hz). **(B)** Basal synaptic transmission was significantly weaker in Arc^−/−^ KO (blue circles) mice compared to WT (black circles) littermates. **(C)** The magnitude of LTP over the first 5 min post-stimulation (1), and over the last 5 minutes of recording (2), was significantly greater in the Arc^−/−^ KO (blue circles) mice compared to WT (black circles) littermates when LTP was induced using a theta-burst stimulation protocol (4 trains at 10 second inter-burst interval. Each train consisted of 10 bursts at 5Hz, with each burst containing 4 stimuli at 100Hz). Data are represented as mean ± SEM. Scale bars, 0.5 mV (horizontal) and 5 ms (vertical).

### CA1 LTP induced by theta burst stimulation is normal in conditional Arc KO mice

The enhanced magnitude of L-LTP after TBS in the full Arc KO lines may be the result of compensatory mechanisms during development. To temporally restrict the loss of Arc, we used a floxed Arc KO line (Arc^fl/fl^) (Chen et al., 2018) injected at 4 – 6 weeks of age with AAV:CaMKII-GFP-cre (GFP-cre) to knock-out Arc in excitatory neurons of CA1, or AAV:CaMKII-GFP alone (GFP) as a matched injection control. Two weeks after viral expression (6 – 8 weeks of age), slices were prepared for electrophysiology. Slices were used for recordings if GFP was identified along the length of the CA1 region of the hippocampus (Fig. 3 *inset*). To determine the magnitude of Arc depletion, we measured Arc protein levels by western blot analysis from the stimulated CA1 region of a subset of slices, post-recording. Arc was significantly reduced in slices from Arc^fl/fl^ animals injected with GFP-cre compared with GFP alone (GFP 1 ± 0.17, n = 5; GFP-Cre 0.24 ± 0.11, *n* = 5; *t*-test*: p* = 0.006) (Fig. 3A).

**Figure. 3.**
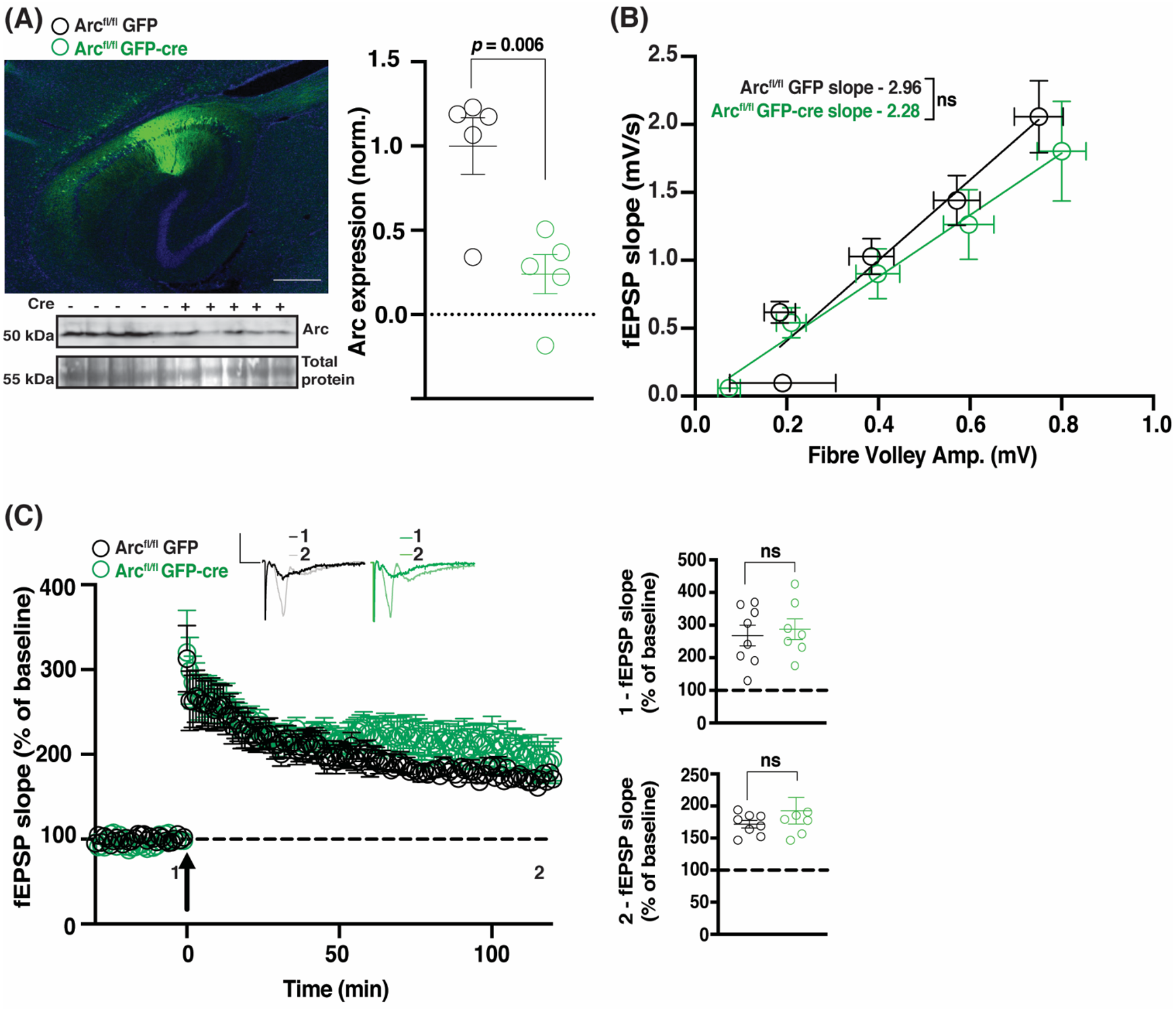
TBS CA1 LTP is normal in Arc cKO animals. **(A)** Arc protein expression was significantly reduced in Arc^fl/fl^ animals injected with GFP-cre (green circles) compared to GFP alone (black circles). **(B)** There was no significant difference in basal synaptic transmission between Arc^fl/fl^ animals injected with GFP and GFP-Cre injected animals. **(C)** There was no significant difference in the magnitude of LTP over the first 5 minutes post-stimulation (1), nor over the last 5 minutes of recording (2), between Arc^fl/fl^ animals injected with GFP-cre and those injected with GFP alone, when LTP was induced using a theta-burst stimulation protocol (4 trains at 10 second inter-burst interval. Each train consisted of 10 bursts at 5Hz, with each burst containing 4 stimuli at 100Hz). Data are represented as mean ± SEM. Scale bars, 0.5 mV (horizontal) and 5 ms (vertical). ***Inset*** – Representative image of GFP expression (green is GFP, blue is DAPI), 2 weeks post-injection of AAV:CaMKII-GFP-cre into the CA1 region of the hippocampus scale bar = 500 μm

**Figure. 4.**
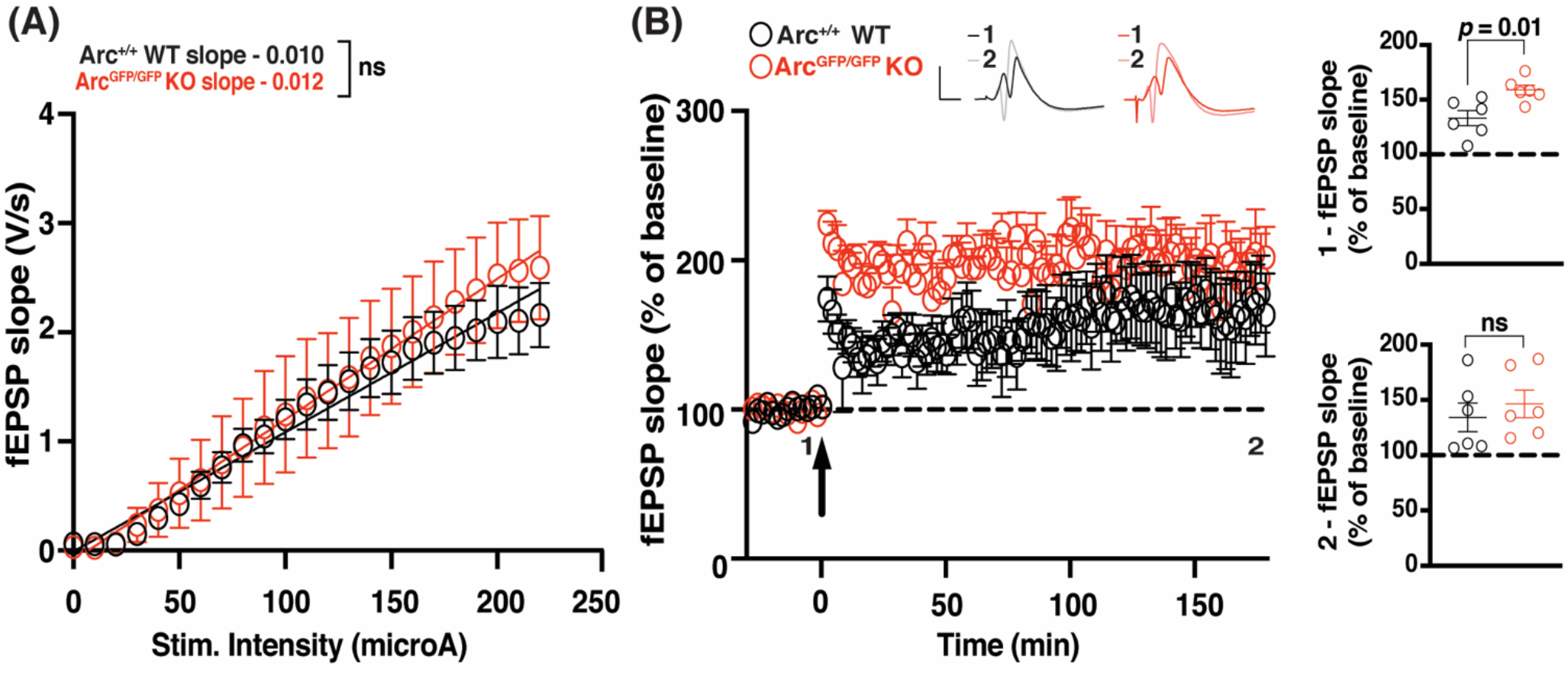
The maintenance of *in vivo* LTP in the dentate gyrus is normal in Arc KO animals. **(A)** There was no significant difference in basal synaptic transmission between the Arc^GFP/GFP^ KO (red circles) and WT (black circles) littermates in the dentate gyrus *in vivo.* **(B)** The magnitude of LTP over the first 5 min post-stimulation (1) was significantly enhanced in Arc^GFP/GFP^ KO animals compared to WT littermates, but was not significantly different over the last 5 minutes of recording (2), when LTP was induced using a theta-burst stimulation protocol (4 trains at 10 second inter-burst interval. Each train consisted of 10 bursts at 5Hz, with each burst containing 4 stimuli at 100Hz). Data are represented as mean ± SEM. Scale bars, 5 mV (horizontal) and 5 ms (vertical).

Basal transmission was similar between Arc^fl/fl^ animals injected with GFP and GFP-Cre injected animals (GFP slope = 2.96 ± 0.69, n = 8; GFP-cre slope = 2.28 ± 0.75, n = 7; linear regression, extra-sum-of-squares F test between curves: F_(1,71)_ = 1.866, *p* = 0.18) (Fig. 3B). Using TBS, we found no difference in the initial magnitude of LTP between Arc^fl/fl^ animals injected with GFP-cre and those injected with GFP alone (GFP 271.1% ± 25.79, n = 8; GFP-Cre 281.1% ± 37.35, n = 7; *t*-test: *p* = 0.68). There was also no significant difference in the magnitude of LTP 2 hours post-induction (GFP 180.8% ± 20.79, n = 8; GFP-Cre 193.4% ± 36.77, n = 7; *t*-test: *p* = 0.32). These results suggest that knock down of Arc post-development has no effect on the induction or maintenance of TBS L-LTP.

### Arc is not required for the maintenance of L-LTP induced by theta burst stimulation in the intact animal

Finally, we assessed whether Arc is required for L-LTP *in vivo* in the dentate gyrus (DG) at medial perforant path-granule cell synapses. There was no significant difference in basal transmission in the dentate gyrus, *in vivo,* between Arc^GFP/GFP^ KO and WT littermates (WT slope = 0.011 ± 0.001, n = 6; Arc^tm1Stl^ KO slope = 0.013 ± 0.002, n = 6; linear regression, extra-sum-of-squares F test between curves: F_(2,272)_ = 2.024, *p* = 0.13). The initial magnitude of LTP induced by TBS was significantly enhanced in Arc^GFP/GFP^ KO animals compared to WT littermate controls (WT 133.13% ± 6.87, n = 6; KO 158.88% ± 4.39, n = 6; *t*-test: *p* = 0.01). However, there was no difference in the magnitude of LTP 3 hours post-stimulation (WT 134.25% ± 13.03, n = 6; KO 146.45% ± 12.51, n = 6; *p* = 0.51). These data further suggest that Arc is not required for the maintenance of L-LTP.

## Discussion

The mechanisms of LTP induction and expression are well defined, but less is known about maintenance. Arc is one of the most robustly expressed genes after LTP induction, with the prevailing view that Arc protein is important for some aspect of LTP (Link et al., 1995; Lyford et al., 1995; Steward et al., 1998; Steward and Worley, 2001). Indeed, Arc protein is increased after LTP induction (Lyford et al., 1995; Steward et al., 1998; Ying et al., 2002; Moga et al., 2004) and disruption of Arc protein expression also results, in some cases, in a deficit in LTP maintenance (Guzowski et al., 2000; Messaoudi et al., 2007). However, relatively few studies have comprehensively tested the role of Arc in hippocampal LTP. Here, we show that LTP is unaffected in both the CA1 and DG of the hippocampus, in multiple Arc KO mouse lines, using an array of LTP induction paradigms. These data indicate that Arc is not essential for the maintenance of LTP.

### Arc is not required for the maintenance of L-LTP in hippocampal slices

Standard NMDAR-dependent LTP in the CA1 has not previously been investigated thoroughly in Arc KO mice. A previous study, using Arc^−/−^ mice, found that LTP was disrupted in CA1 using a patch-clamp induction protocol (depolarization of the postsynaptic cell paired with 200 presynaptic pulses at 1.5 Hz) (Plath et al., 2006). However, we found no significant difference in the maintenance of LTP induced using a standard extracellular field recording HFS protocol (2 × 100 pulses at 100 Hz) (Reymann et al., 1985) in Arc^GFP/GFP^ mice. An NMDAR-independent LTP that requires Type 1 mGluRs receptors has also been shown to be impaired in Arc KO animals (Wang et al., 2016). The induction protocol used for this form of LTP relies upon blocking of NMDAR during a 200 Hz stimulation protocol and It is unclear how this form of LTP relates to memory consolidation processes or if this plasticity is relevant *in vivo*. TBS can also induce robust, persistent L-LTP and is arguably more physiologically relevant than HFS (Larson et al., 1986; Thomas et al., 1998). Surprisingly, we found that L-LTP persisted at a significantly higher magnitude in Arc^GFP/GFP^ KO animals, compared with WT littermates. Recent studies have found behavioral differences between the Arc^−/−^ and the Arc^GFP/GFP^ animal lines (Manago et al., 2016; Gao et al., 2019). This may be due to a neomycin cassette present in Arc^GFP/GFP^, which may lead to side effects (Pham et al., 1996; Scacheri et al., 2001; Gao et al., 2019). We found that basal synaptic transmission is decreased in Arc^−/−^ KO animals but normal in Arc^GFP/GFP^ KO animals. Nevertheless, similar to the Arc^GFP/GFP^ line, we find that TBS-induced L-LTP is enhanced in Arc^−/−^ KO mice. However, TBS induced L-LTP was unaffected in Arc cKO animals, suggesting that this enhanced L-LTP may be due to compensation that arises during development in Arc germline KO animals. Taken together our data suggests that Arc is not required for the maintenance of L-LTP in hippocampal slices.

Since L-LTP maintenance was not impaired in Arc KO mice, we wondered whether the threshold to induce LTP is altered. Metaplasticity, or the plasticity of plasticity, describes the phenomenon where alterations to the basal state of neurons changes the type of plasticity induced by a given specific stimulation paradigm (Abraham and Bear, 1996). For example, the synaptic modification rules based on stimulation frequency, as determined by the Bienstock-Cooper-Munro (BCM) model, can be altered by differences in NMDAR composition at stimulated synapses (Bienenstock et al., 1982; Dudek and Bear, 1992; Jedlicka, 2002; Cooper and Bear, 2012). Thus, we tested whether there was any difference in LTP induced using a protocol that would normally induce only E-LTP (Frey and Morris, 1997) or a moderate frequency (10 Hz) that normally induces little to no change in synaptic strength (Dudek and Bear, 1992). We found no significant difference between genotypes using these induction protocols. Taken together our data suggests that Arc is not required for the induction or maintenance of CA1 LTP.

### Arc does not regulate LTP *in vivo*

Previous studies, using antisense oligodeoxynucleotides (ODN), found that Arc was required for L-LTP in the DG *in vivo* (Guzowski et al., 2000; Messaoudi et al., 2007). We investigated the role of Arc in L-LTP *in vivo* in the DG of Arc^GFP/GFP^ KO animals. We found that the magnitude of L-LTP induction was significantly greater in the first 5 minutes post-stimulation but, by 2 hours, there was no significant difference between KO and WT animals and that it was maintained until at least 3 hours after induction in both genotypes. Interestingly, Plath et al., (2006) found that the magnitude of LTP induction was initially enhanced in Arc^−/−^, but the maintenance was significantly attenuated (Plath et al., 2006). Our experiments described here used the same induction protocol in the DG, but were undertaken in isoflurane-anesthetized mice, where the previous study used urethane anesthesia, and used Arc^GFP/GFP^ rather than Arc^−/−^ mice (Plath et al., 2006). Thus, possible explanations for the difference in the results of the two studies are that maintenance of L-LTP in the DG is differentially affected by either anesthetic regime or Arc knockout strategy. A different study infused ODNs to target Arc into the DG 1.5 hours prior to LTP induction, which had no effect on the initial LTP magnitude but by 5 days post-stimulation LTP magnitude was significantly attenuated (Guzowski et al., 2000). A further study found that when similar ODNs were injected 5 minutes prior to LTP induction or 15 minutes post-LTP induction, synaptic strength transiently decreased to near baseline for around 3 hours, before it recovered to normal levels of potentiation (Messaoudi et al., 2007). Similarly, when ODNs were injected 2 hours post-LTP induction, synaptic strength also decreased to near baseline, but recordings were not made 3 hours later to see if potentiation recovered (Messaoudi et al., 2007). These ODN experiments suggest that acute block of Arc translation may affect expression of LTP but not maintenance. While L-LTP was not affected in our experiments that recorded for hours, we cannot rule out that Arc is involved in the maintenance of potentiation over days.

### How does Arc mediate long-term memory?

Though LTP has long been associated with long-term memory, and long-term memory is critically dependent on Arc expression (Guzowski et al., 2000; Plath et al., 2006; Ploski et al., 2008), the potentiation of synapses is not the only process that underlies the formation and maintenance of long-term memory (Zhang and Linden, 2003; Mozzachiodi and Byrne, 2010; Kyrke-Smith and Williams, 2018; Lisman et al., 2018; Abraham et al., 2019). Similarly, in response to stimulation paradigms that induce LTP, there are widespread changes aside from LTP, such as heterosynaptic depression, changes to intrinsic excitability and the regulation of gene expression, protein synthesis and signalling pathways (Lynch et al., 1977; Andersen et al., 1980; Abraham and Goddard, 1983; Lopez de Armentia et al., 2007; Caroni et al., 2014; Kyrke-Smith and Williams, 2018). We posit that Arc expression driven by LTP induction may contribute to non-LTP plasticity that serves a homeostatic function to maintain overall network stability such as heterosynaptic LTD or synaptic downscaling. A loss of these forms of synaptic weakening in the germline Arc KO mice could account for the exaggerated LTP induction that we and others (Plath et al., 2006) have observed. This idea is also consistent with the finding that Arc interacts with CaMKIIβ at inactive synapses to remove AMPARs and decrease synaptic strength (Okuno et al., 2012; El-Boustani et al., 2018). Similarly, Arc may eliminate small mushroom spines after learning (Nakayama et al., 2015), although it is unclear if this elimination of mushroom spines occurs through heterosynaptic LTD. The process of decreasing the strength of some synapses to enhance the signal to noise ratio of the potentiated synapses, and to maintain a normal dynamic range of plasticity and neuronal activity, may also be critical for long-term memory. Similarly, Arc-dependent homeostatic scaling (Shepherd et al., 2006; Korb et al., 2013) may also be important for stabilizing memory engrams.

Epigenetic mechanisms are also important for long-term memory and L-LTP (Korzus et al., 2004; Levenson et al., 2004; Gräff et al., 2011, 2014; Jarome and Lubin, 2014). Interestingly, Arc has been shown to localize in the nucleus and interact with histone acetyltransferase Tip60, leading to increased acH4K12 (Wee et al., 2014), which is associated with increased gene expression. Arc is also expressed in promyelocytic leukemia nuclear bodies in the nucleus, where it may regulate the expression of plasticity related genes (Korb et al., 2013). We also recently showed that Arc can form virus-like capsids that can transfer RNA cell-to-cell (Pastuzyn et al., 2018). While the role of intercellular Arc in plasticity and memory remains to be determined, it is another mechanism that may underlie long-term memory independent of LTP. Thus, many possibilities exist to explain how Arc can play a critical role in long-term memory without being necessary for L-LTP maintenance. Determining the precise molecular processes governed by Arc will likely provide great insight into the consolidation of memory.

## Author Contributions

All authors contributed to the design of the research. M.K.S, L.J.V and S.F.C performed the research and analysed the data. M.K.S and J.D.S wrote the paper.

## Acknowledgements

We thank Dr. Richard Palmiter (University of Washington) for the generous gift of conditional Arc KO mice and Dr. Paul Worley (The Johns Hopkins University) for the Arc^−/−^ line. This work was funded by the Howard Hughes Medical Institute (M.F.B and R.F.H), and the NIH (J.D.S: R01MH112766).

